# Associations Between the IL-23/IL-17A Cytokine Axis and Ambulatory Blood Pressure in Older Adults

**DOI:** 10.64898/2026.08.06.743406

**Authors:** Benjamin Le Gac, Éric Mukunku Katuvuidi, Adrian Noriega De La Colina, Atef Badji, Maxime Lamarre-Cliché, Diane Vallerand, Hélène Girouard

## Abstract

**Background:** Hypertension, the persistent elevation of blood pressure (BP), is characterized by chronic low-grade inflammation and systemic cytokine release. Circulating cytokines contribute to the development of hypertension and end-organ damage. However, the specific immune profile associated with the progression of hypertension remains unclear. We hypothesize that a plasma cytokine signature reflects early BP changes in older adults.

**Methods:** Seventy participants aged 57-81 years were categorized as normotensive (n = 17), elevated BP (n = 10), or hypertensive (n = 43) based on 24-hour ambulatory BP monitoring and antihypertensive treatment status. Plasma IL-1β, IL-6, IL-10, IL-17A, IL-21, IL-22, IL-23, and TNF-α were quantified using immunoassays. Partial Pearson correlations adjusted for demographic and biochemical covariates were used to assess associations between cytokines, BP, and cytokine-cytokine networks.

**Results:** In untreated hypertensive individuals, plasma IL-23 was positively correlated with 24-hour diastolic BP. Antihypertensive treatment was associated with reduced IL-17A concentrations, which are negatively associated with 24-hour systolic BP. In the elevated BP group, IL-21 concentrations were higher than in normotensive individuals.

To further characterize the cytokine signature, cytokine-cytokine correlations were examined. IL-23 and IL-17A were positively correlated with most interleukins, whereas TNF-α showed few associations. IL-1β exhibited strong correlations with both IL-23 and IL-17A, particularly in untreated participants.

**Conclusion:** IL-23 and IL-17A are associated with BP status and are broadly interconnected with other inflammatory cytokines, highlighting the potential importance of the IL-23/IL-17A axis in the hypertension of development. Early alterations in IL-21 in elevated BP may reflect immune changes that precede the onset of hypertension.

## Introduction

Hypertension, defined as chronically elevated blood pressure (BP), is a leading modifiable risk factor for cardiovascular diseases, cognitive decline, and mortality^1–5^. Although elevated BP is the hallmark of this condition, hypertension is increasingly recognized as a chronic low-grade inflammatory condition^6,7^. Accordingly, the immune system is now known to play a key role in the development and progression of hypertension contributing to end-organ damage^8–12^. The deleterious consequences of hypertension are only partially prevented by antihypertensive treatments, highlighting the contribution of mechanisms beyond BP elevation alone^13,14^. Together, these observations suggest that additional processes, including inflammation, may contribute to long-term vascular and cerebral damage despite adequate BP control.

Inflammation is associated with the systemic release of cytokines by both the innate and adaptive immune systems. Among the inflammatory pathways implicated in hypertension, the interleukin (IL)-23/IL-17A axis has emerged as a key contributor over the past decade^15^. The signature effector cytokine of this axis is IL-17A, primarily produced by γδT cells^16,17^ and T helper (Th)17 cells^15^. Consistent with the activation of this pathway, patients with higher BP exhibit increased circulating Th17 cells^18,19^ and IL-17A levels^18,20–22^.

Th17 cells differentiate from naïve CD4^+^ T cells upon exposure to cytokines such as IL-1β, IL-6, IL-21, and IL-23 released by antigen-presenting cells^23,24^. Among these upstream cytokines, IL-23, IL-1β and IL-6 are increasingly recognized as essential mediators of hypertension pathophysiology^25–35^ and promote IL-17A production^29,36^. Accordingly, IL-1β and IL-6 concentrations are elevated in the serum of hypertensive patients^37,38^. In addition to promoting Th17 responses, polarized T cells can also release IL-21 and IL-22, two other pro-inflammatory cytokines that have been less extensively studied but are increasingly recognized for their involvement in hypertension^39,40^. Conversely, IL-10 acts as a protective anti-inflammatory cytokine in hypertension ^41–45^. Activated T cells also produce IFN-γ and TNF-α, further contributing to the low-grade inflammatory milieu associated with hypertension^31,46,47^.

Interestingly, the IL-23/IL-17A pathway is also a key driver of the pathogenesis of psoriasis^48–50^, a disease in which hypertension is highly prevalent^51^. This clinical association further supports the concept that dysregulation of the IL-23/IL-17A axis may represent a shared inflammatory mechanism linking chronic immune activation to hypertension.

Despite growing evidence linking inflammation to hypertension, important gaps in our understanding remain. One key question is whether a subclinical inflammatory state is already present during the early stages of BP elevation, before the onset of established hypertension. Individuals with elevated BP^52,53^, formerly referred to as prehypertension^54^, represent a particularly relevant population in which to investigate this question. This transitional stage, characterized by BP levels above the normotensive range but below the hypertension threshold, is associated with an increased risk of developing hypertension and cardiovascular diseases^55–57^, and is recognized in current guidelines as a target for early preventive interventions^52,53^.

Determining whether circulating inflammatory markers are altered during this stage of BP elevation could provide valuable insight into the earliest biological changes underlying hypertension and help identify biomarkers of vascular risk before the establishment of overt hypertension. Beyond these early inflammatory changes, the specific cytokine signature most strongly associated with BP elevation in older adults remains incompletely characterized. Moreover, because antihypertensive therapies can themselves modulate circulating cytokine concentrations, distinguishing inflammatory profiles according to treatment status is essential for accurately identifying immune alterations associated with BP elevation. Addressing these knowledge gaps may improve our understanding of the inflammatory mechanisms involved in hypertension development and facilitate the identification of novel biomarkers and therapeutic targets.

We hypothesize that subclinical inflammation, reflected by alterations in circulating cytokine levels, is detectable during the early stages of BP elevation and may serve as an early biomarker of hypertension progression and vascular risk. Considering the growing recognition of the IL-23/IL-17A signaling axis and the role of T cells in hypertension, we selected IL-1β, IL-6, IL-10, IL-17A, IL-21, IL-22, IL-23, and TNF-α as our representative panel of cytokines. We assessed the systemic levels of these cytokines in healthy controls, individuals with elevated BP, and hypertensive participants, including individuals receiving antihypertensive treatment, and evaluated the associations between serum cytokine levels and BP, as well as cytokine-cytokine levels associations in a cohort of older adults.

## Materials and methods

### Data Availability Statement

The anonymized datasets and analytical code supporting the findings of this study have been deposited in a private repository for peer review and will be made publicly available upon publication.

### Study participants

Eighty-four older adults aged 57 to 81 years were recruited from the Research Centre of the *Institut Universitaire de Gériatrie de Montréal* (CRIUGM) participant database, and their eligibility was assessed during a telephone-based screening interview. The ethics review board of CRIUGM and the Montreal Clinical Research Institute approved the study protocol and validated the informed consent documents signed by all participants. The participants underwent a clinical and biological evaluation conducted by a physician (M. Lamarre-Cliche, MD). Exclusion criteria included malignant hypertension (i.e., >180/120 mmHg), diabetes mellitus, heart failure (New York Heart Association class III-IV), myocardial infarction within the previous 3 months, cardiac arrhythmia, rheumatic mitral valve disease, liver failure, renal failure (creatinine clearance < 30 mL/min), stroke, non-compensated thyroid disorder, respiratory problems (i.e., asthma, emphysema), current or past alcohol or drug abuse, and smoking. Seventy participants were retained from the eighty-four originally recruited. Data from fourteen participants were excluded from the final analysis; five due to hemolyzed blood samples and nine due to cytokine concentrations outside the predefined standard cubic spline curve range.

According to the AHA/ACC^52^, Hypertension Canada^53,58^ guidelines and clinical recommendations^59^, participants were classified according to their BP levels and medication: normotensives (n = 17) with a 24-hour SBP (24h-SBP) < 120 mmHg and a 24-hour DBP (24h-DBP) < 80 mmHg; individuals with elevated BP (n = 10) defined by a 120 mmHg ≤ 24h-SBP < 130 mmHg and a 24h-DBP < 80 mmHg; and hypertensives (n = 43) with a 24h-SBP > 130 mmHg or a 24h-DBP > 80 mmHg, and/or receiving current antihypertensive treatment. The hypertensive group included newly diagnosed, untreated participants (n = 15) and those receiving antihypertensive pharmacological treatments (n = 28). The distribution of antihypertensive treatment classes among treated participants is presented in Supplementary Figure 1.

### Blood pressure measurements

Participants underwent BP screening using a 24-hour ambulatory BP monitoring (ABPM) device monitor (Model 90207-3Q; Spacelabs Healthcare®, Snoqualmie, WA, USA) according to Hypertension Canada recommendations^53^. The device installed by nurses (M.G. and H.L.) and BP readings were obtained every thirty minutes over 24 hours^58^. A measuring tape was used to measure the participant’s arm circumference to determine whether they required the standard cuff (12 × 23 cm) or the large (16 × 34 cm) one. The ABPM installation procedures included calibration by manually measuring conventional office BP three times with a sphygmomanometer (using Korotkoff phases I and V) after participants rested in a seated position for at least 15 minutes. The calibration process was performed by a trained specialist (A. Noriega de la Colina, MD, PhD). Validation of the ABPM results was performed by an internal medicine specialist (M. Lamarre-Cliche, MD). Any measurements with fewer than 80% valid BP readings were considered insufficient and excluded. The mean 24-hour BP values were retained for analysis. Participants were given a diary to record daytime (awake) and nighttime (sleep) periods, and activities during the 24 hours^60^.

### Determination of cytokines, creatinine, and calcium concentrations

Plasma samples were analyzed for IL-1β, IL-6, IL-10, IL-17A, IL-21, IL-22, IL-23 and Tumor Necrosis Factor (TNF)-*a* using immunoassays with custom Luminex xMAP technology kits (Human High Sensitivity T-Helper Cells and Human Th17 plex kits) and were read on a Luminex TM 200 system (Luminex, Austin, TX, USA) by Eve Technologies Corporation (Calgary, Alberta, Canada). Serum calcium and creatinine levels were measured at the Hôtel-Dieu de Montréal clinical laboratory.

### Statistical analysis

The normal distribution of our cohort was assessed by using the Shapiro-Wilk test. Age and body mass index (BMI) as general participant characteristics, as well as serum calcium and creatinine levels were compared by Kruskal-Wallis tests. Sex distribution was evaluated by a Fisher’s exact test. Subsequent comparisons between groups were carried out by analysis of covariance (ANCOVA), with age, sex and BMI as well as serum calcium and creatinine levels defined as covariates. Associations between plasma cytokine levels and BP measurements were examined using partial Pearson correlations adjusted for age, sex and BMI as well as serum calcium and creatinine levels. Associations among plasma cytokine levels were examined using partial Pearson correlations adjusted for age, sex, BMI, serum calcium and creatinine levels, 24h-SBP, and 24h-DBP. Covariates were selected based on their potential influence on inflammatory markers and BP measurements. The sample size was estimated with an effect size *f* at 0.4, a power of 80%, 3 groups, and 5 covariates. This a priori analysis suggested a minimum sample size of approximately 64 participants. The outcome data are expressed as mean ± standard deviation. All analysis, tables, and figures were made with R Statistical Software version 4.4.0^61^ and RStudio version 2025.09.2+418^62^. Sample size calculations were performed using G*Power software version 3.1.9.7^63^.

## Results

### General characteristics of participants

Seventy participants were included in this study: 17 normotensive participants, 10 participants with elevated BP, and 43 hypertensives participants. General characteristics, biochemical parameters, and BP measurements were evaluated across these three categories and are reported in Table 1. Age was significantly different across BP categories (H(2) = 8.25; p = 0.016), with hypertensive participants being older than normotensive ones (69.3 years vs. 66.0 years; p = 0.026, Tukey’s test). Sex and BMI were similarly distributed across groups. Serum calcium and creatinine levels differed across BP categories (H(2) = 9.84, p = 0.010; H(2) = 10.23, p = 0.006, respectively). Given these findings and prior evidence, age, sex, BMI, serum calcium and creatinine levels were included as covariates in subsequent ANCOVA and partial regression analysis. 24h-SBP increased across the categories (normotensives: 112.65 ± 1.17 mmHg vs. elevated: 123.80 ± 0.85 mmHg vs. hypertensives: 132.72 ± 1.39 mmHg, F(2,62) = 27.04, p = 3.61 x 10^-9^) as did 24h-DBP (normotensives: 68.65 ± 4.01 mmHg vs. elevated: 71.90 ± 4.56 mmHg vs. hypertensives: 78.14 ± 9.95 mmHg, F(2,63) = 8.32, p = 6.31 × 10⁻⁴), with the highest values observed in the hypertensive group.

**Table 1.** General characteristics of normotensive, elevated BP, and hypertensive participants.

|  |  |  |  |  |
| --- | --- | --- | --- | --- |
| Characteristic | Normotensive<br>(n=17) | Elevated<br>(n=10) | Hypertensive<br>(n=43) | Statistic &<br>P-value |
| **General** |  |  |  |  |
| **Age (years) <sup>a</sup>** | 66.0 (3.04) | 67.35 (4.28) | 69.26 (4.64) | H<sub>(2)</sub>=8.25<br>P=0.016* |
| **Sex <sup>b</sup>** | 7 women / 10<br>men | 7 women / 3<br>men | 20 women / 23<br>men | P=0.376 |
| **BMI (kg/m<sup>2</sup>) <sup>a</sup>** | 25.93 (3.18) | 27.92 (4.32) | 27.48 (4.08) | H<sub>(2)</sub>=2.53<br>P=0.282 |
| **Biochemical analysis <sup>a</sup>** |  |  |  |  |
| **Calcium**<br>**(mg/dL)** | 2.31 (0.11) | 2.24 (0.12) | 2.41 (0.34) | H<sub>(2)</sub>=9.84<br>P=0.010** |
| **Creatinine**<br>**(μmol/L)** | 68.82 (19.27) | 65.50<br>(10.71) | 78.58 (18.91) | H<sub>(2)</sub>=10.23<br>P= 0.006** |
| **Blood Pressure measurements <sup>c</sup>** |  |  |  |  |
| **24h SBP**<br>**(mmHg)** | 112.65 (4.83) | 123.80<br>(2.70) | 132.72 (9.08) | $F_{(2,62)}=27.04$<br>$P=3.61 \times 10^{-9***}$ |
| **24h DBP**<br>**(mmHg)** | 68.65 (4.01) | 71.90 (4.56) | 78.14 (9.95) | $F_{(2,62)}=8.32$<br>$P=6.31 \times 10^{-4***}$ |
| Abbreviations: BMI, Body mass index; SBP, systolic blood pressure; DBP, diastolic blood pressure |  |  |  |  |
| <sup>a</sup>Kruskal-Wallis test; <sup>b</sup>Fischer's exact test; <sup>c</sup>ANCOVA, controlling for age, sex, BMI, and serum calcium and creatinine levels. Mean (SD). *p < 0,05; **p < 0,01, ***p<0.001 |  |  |  |  |

### Plasma cytokine levels across blood pressure categories

Plasma cytokine levels were measured to assess the relationship between BP and systemic inflammation. We first examined plasma cytokine concentrations across the three BP categories. Although no statistically significant differences were detected, several cytokines showed consistent trends across groups.

Plasma IL-21 concentrations appeared higher in the elevated BP category; however, differences across BP categories did not reach statistical significance (Table 2, F(1,63)=2.87, p = 0.064). Plasma levels of IL-1β, IL-6, IL-10, IL-22, IL-23, and TNF-α remained unchanged across BP categories (Table 2), including among participants who were untreated (Supplementary Table 1). Plasma IL-17A levels likewise remained unchanged in the overall cohort. Notably, the hypertensive group included participants receiving antihypertensive treatments, which are known to influence circulating cytokine profiles^64–66^. In our dataset, plasma IL-17A levels were significantly lower in treated than in untreated hypertensive participants (2.36 ± 0.53 vs. 8.14 ± 2.53 pg/mL; F(1,63) = 7.03, p = 0.010, Table 3). Other cytokines did not show significant treatment-related differences, although decreasing trends were observed for IL-1β, IL-6, IL-21, IL-22, and IL-23, while TNF-α showed a tendency to increase with antihypertensive therapy.

**Table 2.**
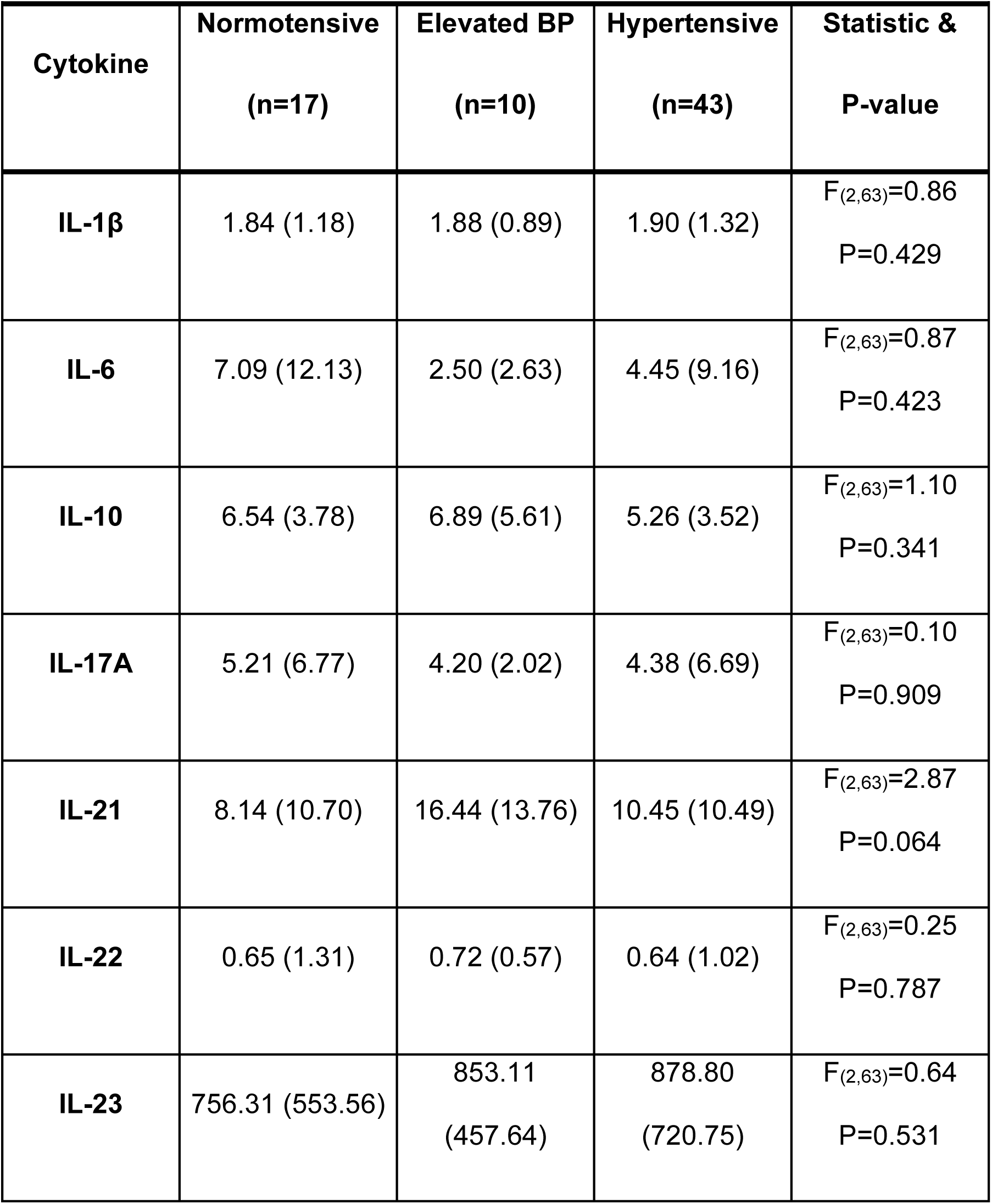
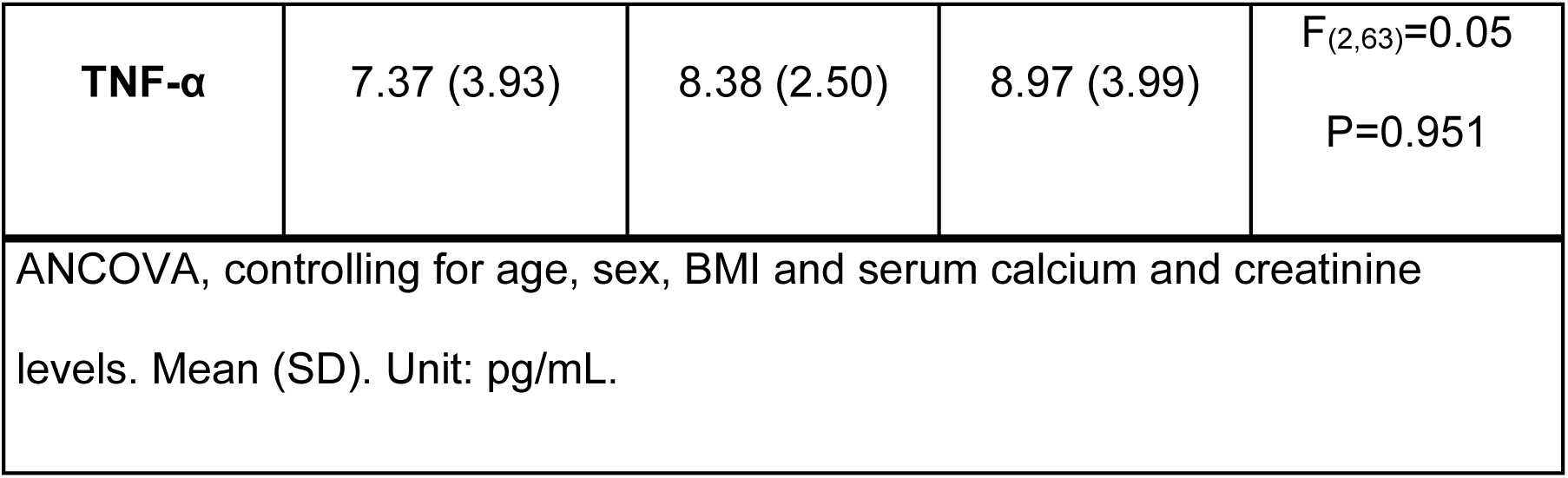
Plasma cytokine levels in normotensive, elevated BP and hypertensive participants.

| **Cytokine** | **Normotensive**<br>**(n=17)** | **Elevated BP**<br>**(n=10)** | **Hypertensive**<br>**(n=43)** | **Statistic &**<br>**P-value** |
| --- | --- | --- | --- | --- |
| **IL-1β** | 1.84 (1.18) | 1.88 (0.89) | 1.90 (1.32) | F<sub>(2,63)</sub>=0.86<br>P=0.429 |
| **IL-6** | 7.09 (12.13) | 2.50 (2.63) | 4.45 (9.16) | F<sub>(2,63)</sub>=0.87<br>P=0.423 |
| **IL-10** | 6.54 (3.78) | 6.89 (5.61) | 5.26 (3.52) | F<sub>(2,63)</sub>=1.10<br>P=0.341 |
| **IL-17A** | 5.21 (6.77) | 4.20 (2.02) | 4.38 (6.69) | F<sub>(2,63)</sub>=0.10<br>P=0.909 |
| **IL-21** | 8.14 (10.70) | 16.44 (13.76) | 10.45 (10.49) | F<sub>(2,63)</sub>=2.87<br>P=0.064 |
| **IL-22** | 0.65 (1.31) | 0.72 (0.57) | 0.64 (1.02) | F<sub>(2,63)</sub>=0.25<br>P=0.787 |
| **IL-23** | 756.31 (553.56) | 853.11<br>(457.64) | 878.80<br>(720.75) | F<sub>(2,63)</sub>=0.64<br>P=0.531 |
| **TNF-α** | 7.37 (3.93) | 8.38 (2.50) | 8.97 (3.99) | $F_{(2,63)}=0.05$<br>$P=0.951$ |
| ANCOVA, controlling for age, sex, BMI and serum calcium and creatinine levels. Mean (SD). Unit: pg/mL. |  |  |  |  |

**Table 3.** Plasma cytokine levels in hypertensive participants according to their treatment.

| Cytokine | Untreated (n=15) | Treated (n=28) | Statistic & P-value |
| --- | --- | --- | --- |
| **IL-1β** | 1.98 (1.72) | 1.86 (1.08) | $F_{(1,63)}=0.022$<br>P=0.883 |
| **IL-6** | 8.04 (14.37) | 2.53 (3.54) | $F_{(1,63)}=2.279$<br>P=0.140 |
| **IL-10** | 5.16 (3.79) | 5.31 (3.44) | $F_{(1,63)}=0.104$<br>P=0.749 |
| **IL-17A** | 8.14 (9.78) | 2.36 (2.82) | **$F_{(1,63)}=7.305$**<br>**P=0.010\*** |
| **IL-21** | 14.72 (14.09) | 8.17 (7.26) | $F_{(1,63)}=4.033$<br>P=0.052 |
| **IL-22** | 0.91 (1.13) | 0.50 (0.94) | $F_{(1,63)}=0.235$<br>P=0.630 |
| **IL-23** | 1022.19 (838.42) | 801.98 (652.72) | $F_{(1,63)}=0.107$<br>P=0.745 |
| **TNF-α** | 8.52 (3.66) | 9.22 (4.20) | $F_{(1,63)}=0.510$ |
|  |  |  | P=0.480 |
| ANCOVA, controlling for age, sex, BMI and serum calcium and creatinine level. Mean (SD). Unit: pg/mL. *p < 0,05 |  |  |  |

Taken together, category-based analysis suggested heterogeneous cytokine patterns across BP groups, including numerically higher IL-21 levels and higher IL-17A levels in untreated hypertensive participants compared with treated hypertensive individuals. Although group differences were not statistically significant, modest numerical shifts, particularly for IL-23, raised the possibility that continuous BP measures might capture associations not evident from categorical comparisons. We therefore next examined associations between cytokine levels and ambulatory BP measurements.

### Associations between plasma cytokine levels and blood pressure

Linear associations between plasma cytokine levels and 24h-SBP or 24h-DBP were assessed by partial Pearson correlations adjusted for age, sex, BMI, serum calcium and creatinine levels (Table 4). Supplementary Figure 2 shows the raw distributions of plasma cytokine levels versus 24h-SBP and 24h-DBP, illustrating the range and variability of the data. Interestingly, IL-23 was significantly and positively correlated with 24h-DBP in the whole cohort (r = 0.323; p = 0.009). This association between IL-23 levels and 24h-DBP persisted when the analysis was restricted to hypertensive participants only (r = 0.347; p = 0.03). As reported in the literature, antihypertensive treatments can modulate plasma cytokine concentrations, thereby influencing their correlation with BP. We next assessed associations stratified by treatment status (Table 5). IL-23 showed the strongest positive correlation with 24h-DBP in the untreated hypertensive group (r = 0.652; p = 0.041). Interestingly, IL-17A was negatively correlated to 24h-SBP among treated participants (r = -0.511; p = 0.013). We did not observe other associations between plasma cytokine levels and BP in this cohort.

**Table 4.** Partial Pearson correlations between cytokine levels and blood pressure.

|  | Whole cohort |  |  |  | Normotensive |  |  |  | Elevated |  |  |  | Hypertensive |  |  |  |
| --- | --- | --- | --- | --- | --- | --- | --- | --- | --- | --- | --- | --- | --- | --- | --- | --- |
| Cytokine | 24h SBP |  | 24h DBP |  | 24h SBP |  | 24h DBP |  | 24h SBP |  | 24h DBP |  | 24h SBP |  | 24h DBP |  |
|  | R | P-value | R | P-value | R | P-value | R | P-value | R | P-value | R | P-value | R | P-value | R | P-value |
| *IL-1β* | 0.200 | 0.111 | 0.126 | 0.319 | 0.147 | 0.650 | -0.315 | 0.319 | 0.738 | 0.154 | -0.314 | 0.607 | 0.191 | 0.251 | 0.081 | 0.627 |
| *IL6* | -0.124 | 0.324 | -0.070 | 0.580 | -0.031 | 0.923 | -0.066 | 0.839 | 0.493 | 0.399 | -0.614 | 0.271 | -0.162 | 0.332 | -0.059 | 0.726 |
| *IL10* | -0.094 | 0.459 | -0.066 | 0.603 | -0.013 | 0.968 | -0.194 | 0.545 | 0.756 | 0.140 | -0.699 | 0.189 | 0.014 | 0.933 | 0.045 | 0.790 |
| *IL17A* | 0.017 | 0.891 | -0.042 | 0.741 | 0.105 | 0.746 | -0.539 | 0.071 | 0.574 | 0.312 | -0.422 | 0.479 | 0.061 | 0.714 | 0.041 | 0.805 |
| *IL21* | 0.079 | 0.531 | 0.069 | 0.586 | -0.197 | 0.539 | 0.073 | 0.822 | -0.342 | 0.573 | 0.527 | 0.361 | -0.034 | 0.839 | 0.042 | 0.803 |
| *IL22* | 0.046 | 0.715 | 0.039 | 0.756 | 0.145 | 0.654 | -0.240 | 0.452 | 0.170 | 0.784 | 0.139 | 0.824 | -0.034 | 0.837 | 0.081 | 0.629 |
| *IL23* | 0.201 | 0.108 | **0.323\*\*** | **0.009\*\*** | -0.098 | 0.762 | -0.170 | 0.597 | -0.090 | 0.886 | 0.429 | 0.471 | 0.237 | 0.152 | **0.347\*** | **0.033\*** |
| *TNF-α* | -0.131 | 0.297 | -0.157 | 0.211 | -0.539 | 0.071 | 0.075 | 0.816 | -0.106 | 0.866 | -0.646 | 0.239 | -0.187 | 0.261 | -0.214 | 0.197 |

Partial Pearson correlation adjusted for age, sex, BMI, serum calcium and creatinine levels. Data are shown as partial R correlation values and p-values. \* p < 0.05; \*\* p <

**Table 5.** Partial Pearson correlation between cytokine levels and blood pressure according to the treatment.

|  | Untreated hypertensive (n=15) |  |  |  | Treated hypertensive (n=28) |  |  |  |
| --- | --- | --- | --- | --- | --- | --- | --- | --- |
|  | 24h SBP |  | 24h DBP |  | 24h SBP |  | 24h DBP |  |
| Cytokine | R | P-value | R | P-value | R | P-value | R | P-value |
| *IL-1β* | 0.584 | 0.076 | 0.152 | 0.675 | -0.162 | 0.461 | 0.010 | 0.963 |
| *IL6* | -0.122 | 0.738 | 0.147 | 0.686 | -0.156 | 0.478 | -0.224 | 0.304 |
| *IL10* | -0.124 | 0.732 | -0.014 | 0.970 | 0.030 | 0.893 | 0.046 | 0.837 |
| *IL17A* | 0.307 | 0.388 | 0.088 | 0.809 | **-0.511\*** | **0.013\*** | -0.276 | 0.203 |
| *IL21* | -0.008 | 0.983 | 0.534 | 0.112 | -0.089 | 0.688 | -0.128 | 0.562 |
| *IL22* | 0.217 | 0.547 | 0.594 | 0.070 | -0.143 | 0.514 | 0.035 | 0.873 |
| *IL23* | 0.065 | 0.858 | **0.652\*** | **0.041\*** | 0.182 | 0.405 | 0.320 | 0.137 |
| *TNF-α* | -0.371 | 0.292 | -0.114 | 0.753 | -0.140 | 0.525 | -0.286 | 0.187 |

Partial Pearson correlation adjusted for age, sex, BMI, serum calcium and creatinine levels.
Data are shown as partial R correlation values and as p-values. \*p < 0.05

Taken together, these findings suggest that IL-23 is associated with diastolic BP across the cohort, with the strongest relationship observed among untreated hypertensive participants. In addition, plasma IL-17A levels were modulated by antihypertensive treatments and were negatively correlated with 24h-SBP in treated participants.

### Associations between plasma cytokine levels

Given the reported involvement of the IL-23/IL-17A axis in hypertension, we next investigated associations among circulating cytokines using partial Pearson correlations adjusted for age, sex, BMI, serum calcium, creatinine, and ambulatory BP. Correlation matrices and raw distributions of cytokine concentrations were generated, illustrating the range and variability of the data. Associations were assessed across all participants (Figure 1A) and untreated participants including normotensives participants, participants with elevated BP and untreated hypertensive individuals (n = 42; Figure 1B) as we previously observed an effect of antihypertensive treatment on IL-17A and IL-23 (Table 3 and 5). Several correlations strengthened in untreated participants, suggesting that antihypertensive treatment attenuates cytokine coupling (Figure 1B). Notably, TNF-α was not considered further in network interpretation owing to weak or absent correlations.

**Figure 1.**
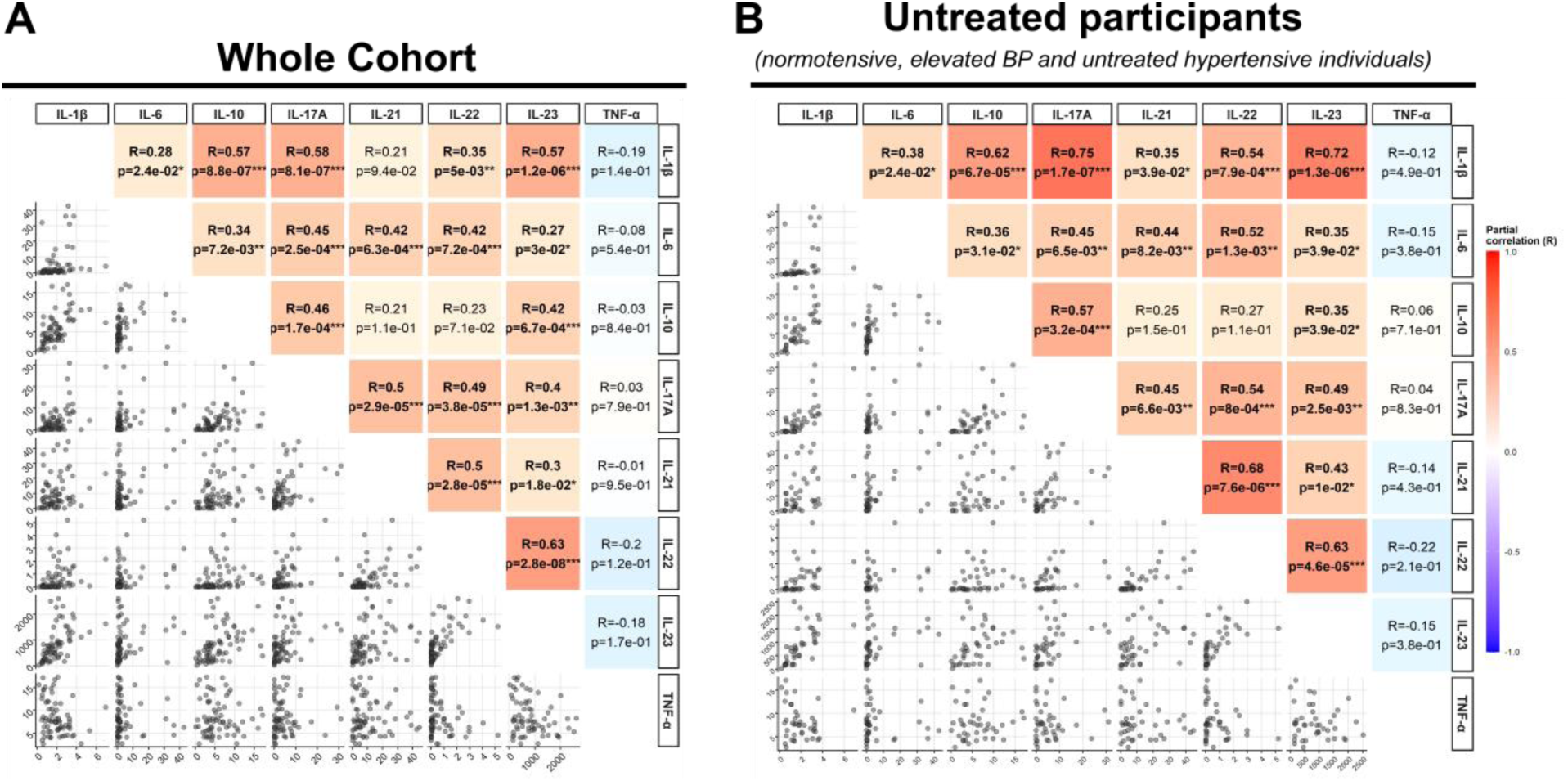
Partial Pearson correlation between cytokine levels. Partial Pearson correlation adjusted for age, sex, BMI, serum calcium and creatinine levels, 24h-SBP and 24h-DBP (A) for the whole cohort (n = 70) or (B) only untreated participants including normotensive, elevated BP and hypertensive individuals (n = 42). The upper part displays the correlation matrix with the partial correlation R value. Colours in the figure represent partial correlation coefficients from − 1 to 1. Significant at p-value *p< 0,05, **p< 0,01 and ***p< 0,001. The lower part displays the raw distribution between cytokines. Each point represents an individual. Unit: pg/mL.

In the whole cohort, IL-23 was significantly and positively correlated with all interleukins (Figure 1A). Specifically, its plasma levels showed moderate correlations with IL-1β (R = 0.57, p = 1.2 x 10^-6^) and IL-22 (R = 0.63, p = 2.8 x 10^-8^) concentrations, which were even stronger in untreated participants. We also observed weak but significant correlations of IL-23 with IL-6 (R = 0.27, p = 0.03), IL-17A (R = 0.4, p = 0.0013) and IL-21 (R = 0.3, p = 0.018). Similarly, plasma IL-17A and IL-1β levels were correlated with several interleukins. IL-17A and IL-1β were positively correlated (R = 0.58, p = 8.1 x 10^-7^), with a stronger relationship in untreated participants (R = 0.75, p = 1.7 x 10^-7^).

Among the remaining cytokines, plasma IL-6 concentrations were overall weakly associated with other interleukins. Plasma levels of the anti-inflammatory cytokine IL-10 were associated with the pro-inflammatory cytokines IL-1β, IL-6, and IL-23, along with IL-17A. IL-21 and IL-22 displayed similar association patterns with IL-6, IL-17A, IL-23 and with each other.

These findings reveal a highly interconnected cytokine correlation network highlighting the central position of the IL-23/IL-17A axis after adjustment for general characteristics and BP. Furthermore, IL-1β emerged as a highly connected node alongside the IL-23/IL-17A axis, particularly in untreated participants.

## DISCUSSION

Our findings suggest that BP elevation in older adults is associated with selective alterations in inflammatory pathways rather than a broad systemic activation. Specifically, plasma IL-23 concentrations showed a gradual increase across normotensive participants, participants with elevated BP, and hypertensive groups in this cohort of older adults. This trend was further supported by a positive association between IL-23 and 24h-DBP, evident across all participants and particularly pronounced among untreated hypertensive participants. In contrast, most circulating cytokines did not differ significantly across BP categories, highlighting the importance of examining continuous BP measures and cytokine relationships beyond categorical comparisons. We also identified a potential modulatory effect of antihypertensive therapy on inflammatory profiles, with lower IL-17A levels in treated hypertensive participants and a negative association between IL-17A and 24h-SBP among treated individuals. Finally, both IL-23 and IL-17A were positively associated with multiple other interleukins. IL-1β was strongly correlated with IL-23 and IL-17A, particularly in untreated participants. Together, these findings support the involvement of an interconnected IL-23/IL17A-centered cytokine network in BP regulation and hypertension-related inflammation in older adults. Additional non-significant changes, including increased IL-21 concentrations in the elevated BP group, suggest that early immune alterations may precede the development of overt hypertension and warrant further investigation.

### Cytokines and BP

Plasma IL-23 and IL-17A levels are up-regulated during hypertension in both observational human studies and mouse models of hypertension, and they play key roles in the pathophysiology of hypertension^15,67^. In our cohort of older adults, plasma IL-23 concentrations were positively correlated with 24h-DBP in all participants, in untreated individuals and in hypertensive participants. These findings are consistent with previous clinical studies reporting a positive association between plasma IL-23 levels and BP elevation. Specifically, IL-23 levels were correlated in hypertensive patients with both 24h-SBP and 24h-DBP^32^ and were elevated in a young obese hypertensive population^33^. Nonetheless, this association between plasma IL-23 concentration and BP could depend on the time since diagnosis. Indeed, IL-23 levels were not associated with 24h-AMBP in a cohort including patients with newly diagnosed hypertension^68^. The duration of hypertension was not assessed in our study.

In our study, plasma IL-17A levels remained unchanged across BP groups. The role of IL-17A is context-dependent and may differ according to the type of hypertension. Higher levels of plasma IL-17A were found in patients with hypertension^18,69,70^ and those with hypertension as a comorbidity and who were affected by type 2 diabetes^20^, placental insufficiency^71^ or HIV^72^. Moreover, a positive linear relationship between circulating IL-17A and mean arterial blood pressure was found in patients with end-stage renal disease on hemodialysis with refractory hypertension^73^. When considering earlier stages of BP elevation, IL-17A has previously been reported as increased in prehypertensive patients^21^. Differences in BP classification and measurement methods may partly explain these discrepancies across studies. In our cohort, BP categories were defined using 24h-ABPM, which provides more reliable BP assessment than office measurements by reducing misclassification related to white-coat and masked hypertension. In addition, previous studies relied on older definition of “prehypertension” from the Seventh Report of the Joint National Committee on Prevention, Detection, Evaluation, and Treatment of High Blood Pressure^54^ whereas we followed AHA/ACC^52^ and Hypertension Canada^53,58^ guidelines and clinical recommendations^59^. Taken together, the observed trends in IL-23 and IL-17A levels may reflect activation of the Th17 axis, which has been repeatedly implicated in vascular inflammation and endothelial dysfunction in hypertension.

Beyond cytokines involved in Th17 differentiation, other inflammatory mediators have also been implicated in hypertension but showed limited associations with BP in our cohort. There is evidence that IL-1β and IL-6 contribute to hypertension. First, they both drive CD4+ T cell polarization and therefore promote IL-17A production. In humans, hypertension is associated with increased plasma levels of IL-6^37,74–76^ and IL-1β^28,77,78^. By contrast, we did not observe any correlation between these cytokines and BP, even when the analysis was restricted to untreated participants. This absence of correlation suggests that, in this older population, systemic IL-1β/IL-6 signaling may not represent the primary driver of the cytokine-BP relationships observed here, or that their effects are temporally confined, compartmentalized, masked by other factors. Nevertheless, the modest reduction in circulating IL-6 concentrations observed in the elevated BP group may reflect transient inflammatory changes during early BP elevation.

The role of IL-21 in hypertension is understudied. Recently, IL-21 was reported to be increased in an angiotensin II-induced mouse model of hypertension, and an anti-IL-21 treatment was shown to lower BP^40^. In addition, the authors demonstrated that CD4+ T cells production of IL-21 correlated with SBP and IL-17A. In our cohort, we observed a non-significant increase in IL-21 in the elevated BP category. Although the increase did not reach statistical significance, these exploratory observations raise the possibility that IL-6 and IL-21 fluctuations may accompany early immune alterations during BP elevation and warrant further longitudinal investigation.

Circulating IL-22, TNF-α, and IL-10 levels showed minimal variation across BP categories and were not correlated with BP in our cohort. Previous studies have reported elevated plasma IL-22 levels in hypertensive patients, with positive correlations with both SBP and DBP^39^. In contrast, the contribution of TNF-α to hypertension pathophysiology remains unclear, as most evidence originates from preclinical studies and has yielded conflicting findings^7,46,79^. IL-10, an anti-inflammatory cytokine, is upregulated in rodent models of hypertension and has been associated with BP reduction and protection against hypertension-induced vascular dysfunction^41,42,44,80^. However, circulating IL-10 levels have been reported to be reduced in untreated hypertensive patients^43^, and remain unchanged in preeclampsia^81^.

Nevertheless, circulating cytokine concentrations may not fully capture the compartmentalized inflammatory signaling that accompanies hypertension. Immune activation within the kidney, vasculature, heart, and perivascular tissues can diverge substantially from circulating cytokine profiles. In humans, IL-17A-positive cells, including Th17 and γδ T lymphocytes, have been identified in kidney biopsies from patients with hypertensive nephrosclerosis^82^. Additional human support comes from asymptomatic hypertensive patients, in whom coronary sinus sampling detected differences in TNF-α and IL-6 concentrations^83^. In parallel, experimental models of hypertension show immune cell accumulation in lymph nodes, aorta, kidneys, and perivascular tissue, together with increased local cytokine signaling in vascular and renal compartments^19,40,84–86^. Even so, plasma cytokine measurements remain clinically useful because they can be obtained by minimally invasive sampling and may still facilitate the development of accessible biomarkers for risk stratification and disease monitoring. Accordingly, the absence of a systemic association for a given cytokine does not exclude a relevant contribution at the tissue level.

Overall, our results underscore a heterogeneous pattern in cytokine-BP associations, with some pathways, such as the IL-23/IL-17A axis, showing stronger relevance than others. Because inflammatory signaling arises from coordinated networks rather than isolated mediators^87,88^, we next examined cytokine-cytokine relationships to further characterize the inflammatory signature associated with BP elevation.

### Cytokine-cytokine interactions in hypertension

Because these cytokines originate from interacting innate and adaptive immune compartments, examining their associations may provide insight into how coordinated inflammatory signaling contributes to hypertension. The IL-23/IL-17A axis has been implicated in the low-grade inflammation associated with hypertension^15,48–50^. In our study, IL-23 was positively associated with IL-17A, IL-21 and IL-22. This finding is consistent with the established role of IL-23 as an upstream regulator of Th17-related responses^24,29,89^. IL-23 and IL-17A have been found to be positively correlated in human plasma^90^. We found that IL-1β, IL-6, and IL-23 were positively associated with IL-17A, consistent with previously described interactions regulating IL-17A responses^24,89^. Interestingly, IL-10 was found to be positively associated with IL-1β, IL-6, IL-17A and IL-23. This anti-inflammatory cytokine is capable of inhibiting the synthesis of some pro-inflammatory cytokines, such as TNF-α and IL-1β^91^ and reducing BP in rodent models^41–45^. These positive associations may reflect a counter-regulatory response in which IL-10 increases alongside pro-inflammatory activation of the IL-23/IL-17A axis. Finally, plasma TNF-α levels showed no significant correlations with the other cytokines, supporting a role outside of the observed immune network in our cohort. Because antihypertensive therapies may themselves modulate cytokine signaling, these interactions should be interpreted within the context of treatment exposure.

### Effect of antihypertensive treatments on cytokine levels

We found that antihypertensive treatments were associated with lower plasma IL-17A levels in hypertensive participants, and IL-17A was negatively correlated with 24-h SBP in treated hypertensive participants. First-line antihypertensive treatments targeting the renin-angiotensin-aldosterone system, including ARBs and ACE inhibitors used by participants in our cohort, can modulate the IL-23/IL-17A immune axis. These agents can reduce plasma IL-17A, IL-6, IL-23 and TNF-α levels while increasing IL-10 concentrations^66,74,92–94^. However, the limited number of treated participants restricts our ability to assess the effect of each antihypertensive class on plasma cytokine levels. A more detailed characterization of treatment-specific immune signatures would help determine whether inflammation is reduced as a direct pharmacological effect, a consequence of improved BP control, or both.

### Cytokines: potential targets or biomarkers for hypertension?

Inflammation contributes to hypertension development, progression, and end-organ damage^7^, and may also contribute to links between hypertension and neurodegenerative diseases^95^. This raises the question of whether inflammatory pathways, and cytokines in particular, could serve as therapeutic targets or reliable biomarkers^50,92,96^. In this study, we assessed cytokine levels in the plasma to characterize this inflammatory component in older adults. Our findings identified IL-23 as a cytokine associated with BP and positioned within an interconnected inflammatory network. These observations raise important questions regarding the therapeutic relevance of inflammatory pathways in hypertension. The IL-23/IL-17A axis has shown some promising therapeutic potential for autoimmune diseases such as Crohn’s disease, psoriasis and multiple sclerosis, notably through the use of monoclonal antibodies^15,50^. IL-23 has also been proposed as a therapeutic target in atherosclerosis^97^. However, interventions aimed at modulating the IL-23/IL-17A axis yield mixed outcomes with limited efficiency or disease worsening in Crohn’s disease^98,99^. In addition, anti-IL-23 treatments are associated with adverse effects that must be carefully considered^100^. Given these uncertainties, targeting the IL-23/IL-17A axis in hypertension should be approached with caution and evaluated thoroughly.

However, this IL-23 and 24h-DBP association may warrant investigation as a candidate biomarker for the early detection of DBP elevation, called isolated diastolic hypertension when above 90 mmHg^101^. Isolated diastolic hypertension accounts for approximately 20% of hypertension cases in adults and is associated with increased cardiovascular risk independently of age and sex^102^. We observed an increase in IL-21 in the elevated BP group in our cohort. These cytokine patterns may represent early immunological signatures associated with future hypertension and cardiovascular risk. Longitudinal studies incorporating baseline cytokine profiles obtained before BP elevation will be important to determine whether these signals precede BP elevation. Individual baseline BP could therefore serve as a personalized control for such analyses.

### Limitations

Several limitations should be considered. The modest sample size may have limited statistical power and constrained the assessment of differential treatment effects. In addition, the cross-sectional design precludes causal inference regarding relationships between cytokine levels and blood pressure. Our analysis was restricted to the cytokines included in the available assay panels, and circulating cytokine levels may not fully reflect tissue-specific inflammatory processes relevant to hypertension. Residual confounding related to antihypertensive treatment cannot be excluded. Moreover, cytokine-cytokine associations should be interpreted cautiously given the number of comparisons performed. Longitudinal studies with repeated blood sampling and ambulatory blood pressure measurements will be required to define cytokine dynamics during BP elevation and treatment. Despite these limitations, the study identifies associations that warrant validation in larger prospective cohorts.

## Conclusion

In conclusion, our study highlights an association between IL-23 levels and BP, with IL-23 occupying a central position within the observed cytokine network. Antihypertensive treatment was also associated with altered IL-17A levels. Although fluctuations IL-21 levels in the elevated BP group were not statistically significant, this trend warrant further investigation. Together, these findings support further investigation of inflammatory cytokine networks, particularly the IL-23/IL-17A axis, during the early stages of hypertension and BP-related vascular risk.

## Abbreviations list

ABPM: Ambulatory blood pressure monitoring
ACC: American college of cardiology
ACEI: Angiotensin converting enzyme inhibitors
AHA: American heart association
ANCOVA: Analysis of covariance
ARB: Angiotensin II receptor blockers
BMI: Body mass index
BP: Blood pressure
CCB: Calcium channels blockers
CD4: Cluster of differentiation 4
CRIUGM: Centre de Recherche de l’Institut Universitaire de Gériatrie de Montréal
DBP: Diastolic blood pressure
IFN-γ: Interferon-gamma
IL: Interleukin
SBP: Systolic blood pressure
Th: T helper
TNF-α: Tumor necrosis factor-alpha

## Acknowledgments

The authors would like to thank nurses Martine Gauthier and Hélène L’Archevêque from the IRCM for their invaluable help with blood sample collection and Eve technologies for their help on cytokines quantification.

## Supplemental Material

**Supplementary Figure 1.**
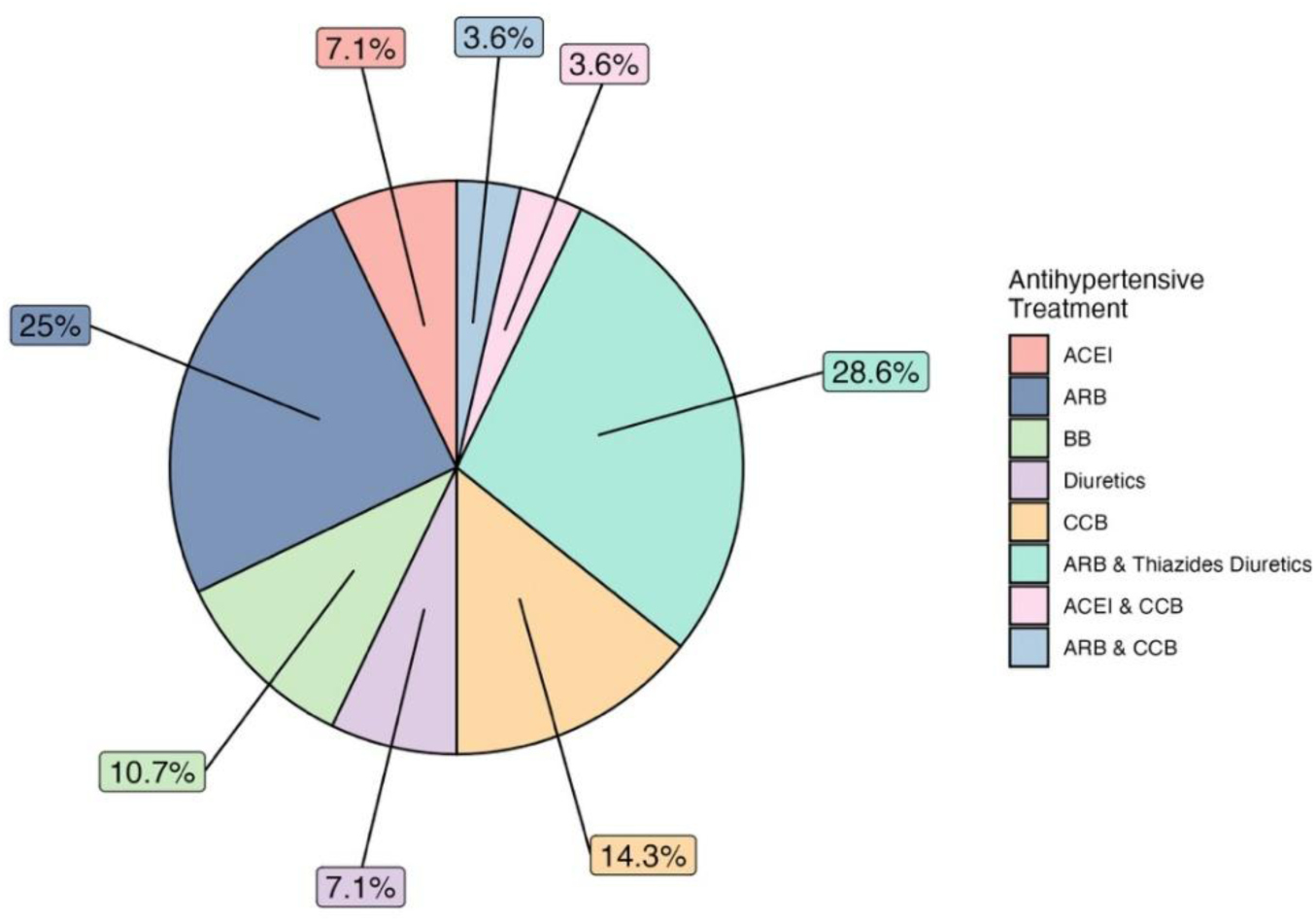
Distribution of antihypertensive treatment classes among treated participants. ACEI: Angiotensin-Converting Enzyme Inhibitors, ARB: Angiotensin II Receptor Blockers, BB: β-Blockers, CCB: Calcium Channel Blockers. Data represented as a percentage of the treated population.

**Supplementary Figure 2.**
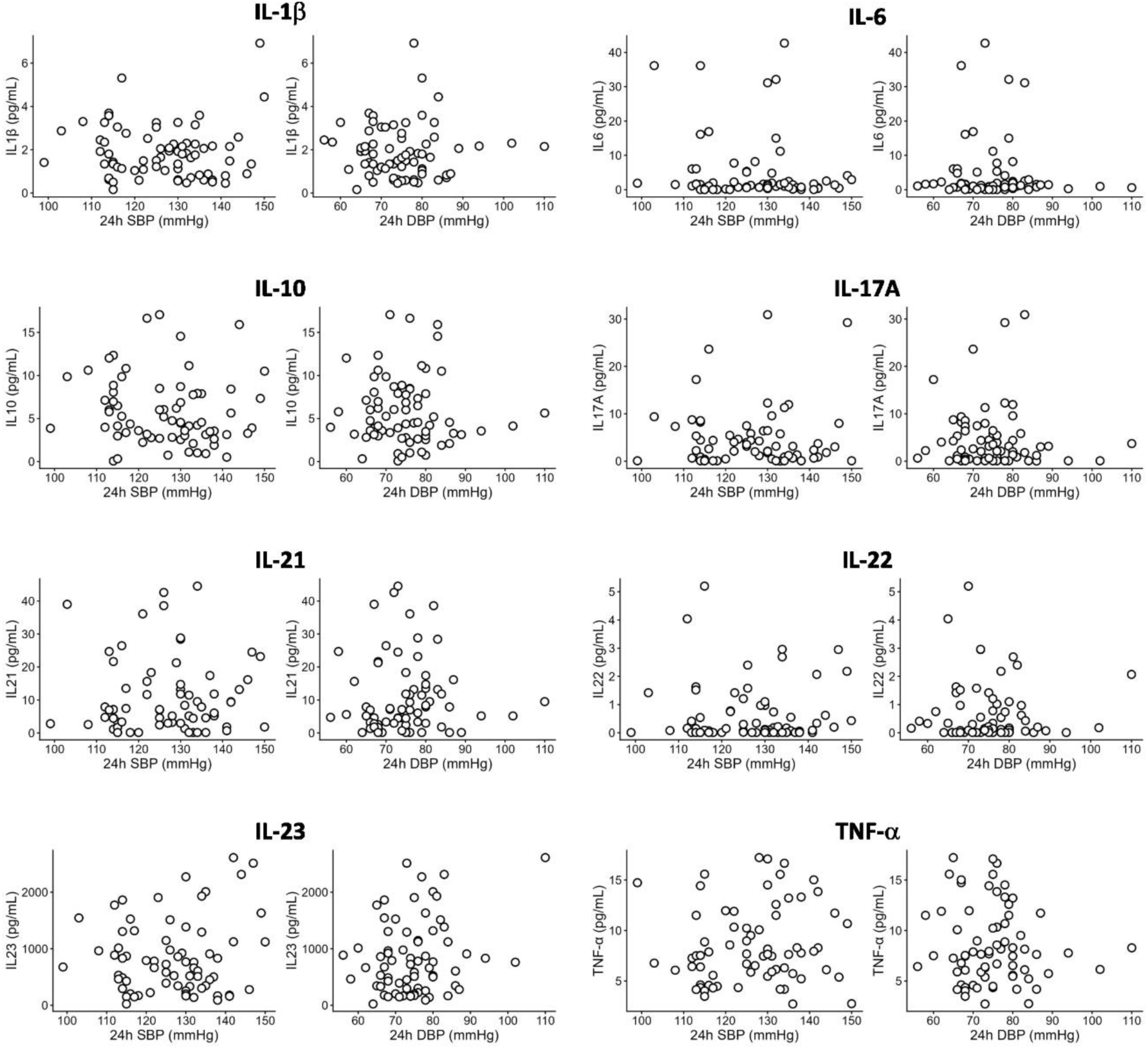
Distribution of cytokine levels according to 24h-SBP and 24h-DBP in all participants. Each point represents the value for one individual.

**Supplementary Table 1.** Plasma cytokine levels in normotensive, elevated BP and untreated hypertensive participants.

| **Cytokine** | **Normotensive**<br>**(n=17)** | **Elevated BP**<br>**(n=10)** | **Untreated**<br>**hypertensive**<br>**(n=15)** | **Statistic &**<br>**P-value** |
| --- | --- | --- | --- | --- |
| **IL-1β** | 1.84 (1.18) | 1.88 (0.89) | 1.98 (1.72) | F<sub>(2,34)</sub>=0.76<br>P=0.476 |
| **IL-6** | 7.09 (12.13) | 2.50 (2.63) | 8,04 (14,37) | F<sub>(2,34)</sub>=1.02<br>P=0.372 |
| **IL-10** | 6.54 (3.78) | 6.89 (5.61) | 5.16 (3.79) | F<sub>(2,34)</sub>=0.33<br>P=0.718 |
| **IL-17A** | 5.21 (6.77) | 4.20 (2.02) | 8,14 (9.78) | F<sub>(2,34)</sub>=1.34<br>P=0.275 |
| **IL-21** | 8.14 (10.70) | 16.44 (13.76) | 14,72 (14,09) | F<sub>(2,34)</sub>=2.63<br>P=0.087 |
| **IL-22** | 0.65 (1.31) | 0.72 (0.57) | 0.91 (1.13) | F<sub>(2,34)</sub>=0.23<br>P=0.797 |
| **IL-23** | 756.31<br>(553.56) | 853.11<br>(457.64) | 1022,19<br>(838,42) | F<sub>(2,34)</sub>=0.87<br>P=0.426 |
| **TNF-α** | 7.37 (3.93) | 8.38 (2.50) | 8.52 (3.66) | F<sub>(2,34)</sub>=0.12<br>P=0.889 |

ANCOVA, controlling for age, sex, BMI and serum calcium and creatinine levels. Mean (SD). Unit: pg/mL.

